# Symptoms of fatigue and depression is reflected in altered default mode network connectivity in multiple sclerosis

**DOI:** 10.1101/505974

**Authors:** Einar August Høgestøl, Gro Owren Nygaard, Dag Alnæs, Mona K. Beyer, Lars T. Westlye, Hanne F. Harbo

**Affiliations:** Department of Neurology, Institute of Clinical Medicine, University of Oslo, Oslo, Norway; Department of Neurology, Oslo University Hospital, Oslo, Norway; KG Jebsen Centre for Psychosis Research, NORMENT, Division of Mental Health and Addiction, Oslo University Hospital, Oslo, Norway & Institute for Clinical Medicine, University of Oslo, Oslo, Norway; Department of Radiology and Nuclear Medicine, Oslo University Hospital, Oslo, Norway; Department of Psychology, University of Oslo, Oslo, Norway

## Abstract

**Background:** Fatigue and depression are frequent and often co-occurring symptoms in multiple sclerosis (MS). Resting-state functional magnetic resonance imaging (rs-fMRI) represents a promising tool for disentangling differential associations between depression and fatigue and brain network function and connectivity. In this study we tested for associations between symptoms of fatigue and depression and DMN connectivity in patients with MS.

**Materials and methods:** Seventy-four MS patients were included on average 14 months after diagnosis. They underwent MRI scanning of the brain including rs-fMRI, and symptoms of fatigue and depression were assessed with Fatigue Severity Scale (FSS) and Beck Depression Inventory II (BDI). A principal component analysis (PCA) on FSS and BDI scores was performed, and the component scores were analysed using linear regression models to test for associations with default mode network (DMN) connectivity.

**Results:** We observed higher DMN connectivity with higher scores on the primary principal component reflecting common symptom burden for fatigue and depression (Cohen’s f^2^=0.075, t=2.17, p=0.03). The secondary principal component reflecting a pattern of low fatigue scores with high scores of depression was associated with lower DMN connectivity (Cohen’s f^2^=0.067, t=-2.1, p=0.04). Using continuous mean scores of FSS we also observed higher DMN connectivity with higher symptom burden (t=3.1, p=0.003), but no significant associations between continuous sum scores of BDI and DMN connectivity (t=0.8, p=0.4).

**Conclusion:** Multivariate decomposition of FSS and BDI data supported both overlapping and unique manifestation of fatigue and depression in MS patients. Rs-fMRI analyses showed that symptoms of fatigue and depression was reflected in altered DMN connectivity, and that higher DMN activity was seen in MS patients with fatigue even with low depression scores.

## Introduction

MS is a heterogeneous disease of the central nervous system (CNS) with typical age of disease onset between 28 and 31 years (1). One of the most common symptoms in multiple sclerosis (MS) is fatigue, affecting up to 90 % of all MS patients (2-4). Fatigue may have a large impact on the daily life of MS patients and may impair both quality of life and ability to work (2-4). Depression is also a common symptom in MS; the lifetime prevalence is reported to be 40-60 % (2, 3, 5). The pathophysiology of these symptoms in MS is not fully understood (2-4, 6).

Magnetic resonance imaging (MRI) is an essential tool in diagnosis and clinical evaluation of MS patients, including follow-up of disease modifying therapies (DMT) (7). Structural MRI studies have shown different patterns of cortical thickness in MS patients who have either fatigue, depression both depression and fatigue. However, these cortical underpinnings only explained 17.3 % of the total variance of the neuropsychiatric symptoms (8). Diverse results are reported concerning the presence and severity of fatigue in relation structural MRI findings in MS (lesions, normal appearing white matter damage or grey matter damage) (3). Some have reported changes in cortico-subcortical pathways such as in the prefrontal cortex, thalamus and basal ganglia in patients with MS-related fatigue (4). Both structural MRI and functional MRI (fMRI) have been applied in many studies with the aim to understand mechanisms responsible for clinical disability, depression, fatigue and cognitive impairment in MS (3).

fMRI studies have shown that the brain is organized in distinct functional networks, and their interplay is central for optimal functioning and health of the brain. Functional connectivity can be conceptualized as the interaction between two different brain regions. Disconnection caused by white matter damage in MS leads to brain network dysfunction, named a disconnection syndrome (3). Both damage to the white and grey matter in MS patients is likely to disrupt brain network connectivity within cortical and sub-cortical networks (9). fMRI has made it possible to assess the integration of activity across distant brain regions and has provided insight into the intrinsic connectivity network. Resting-state (rs) fMRI in MS has mainly been used to study the intrinsic functional architecture and connectivity of the brain and relation to disease progression and clinical impairment (9, 10).

In particular, rs-fMRI has highlighted the role of the default mode network (DMN) as a critical hub for both integration and flow of information (11). The DMN comprises the precuneus, the posterior cingulate cortex (PCC), the angular gyrus, the medial prefrontal cortex (mPFC) and the inferior parietal regions (3, 9). The DMN is most active when a person is not focused on a specific task, often referred to as wakeful rest (11). The DMN has been studied in a wide range of neurological and neuropsychiatric disorders and has provided insights into disease pathophysiology (11). Assuming a role of the DMN in introspection and rumination, DMN changes in MS patients have been proposed to be linked with cognitive dysfunction and depression (11-13). Some fMRI studies have reported cortico-subcortical dysfunction in MS patients with fatigue, also specifically involving fronto-parietal regions and the basal ganglia (3, 4, 14). A recent rs-fMRI study found that specific thalamo-cortical connections explained different components of fatigue in MS patients (14). Thus, there is evidence of altered DMN connectivity in MS patients with symptoms of both depression and fatigue. Although related, these symptoms do not always co-occur, and little is known about the different patterns of DMN alterations with different symptom burden (8). On this background, we aimed to study the common and differential associations between symptoms of fatigue and depression and DMN connectivity using rs-fMRI in MS.

## Materials and methods

### Participants

We included in total 74 MS patients at Oslo University Hospital for a prospective longitudinal study. Some other data from this study have been published earlier (15, 16). All participants were diagnosed between January 2009 and October 2012 with relapsing-remitting MS (RRMS) according to the revised McDonald Criteria (17) and were referred to brain MRI between January 2012 and January 2013. Seven participants did not perform the rs-fMRI sequence, and the remaining 67 participants were used in the current imaging analyses.

Exclusion criteria included age < 18 years or > 50 years, uncertain diagnosis, non-fluency in Norwegian, neurological or psychiatric disease, drug abuse, head trauma, pregnancy and previous adverse gadolinium reaction. The project was approved by the regional ethical committee of South Eastern Norway (REC ID:2011/1846), and all participants received oral and written information and gave their written informed consent.

All participants completed a comprehensive neurological examination, including expanded disability status scale (EDSS) by a Neurostatus certified medical doctor (http://www.neurostatus.net/) and symbol digits modalities test (SDMT) within the same week as their MRI examination. All participants also completed self-reported questionnaires concerning fatigue (Fatigue Severity Scale, FSS; 18), with 9 subscores covering the different dimensions of fatigue, and depressive symptoms (Beck Depressive Inventory II, BDI; 19) with a total of 21 subscores to encompass various features of depression. FSS mean score ≥ 4 was categorized as clinically significant fatigue, while BDI sum score ≥ 14 was categorized as clinically significant depressive symptoms (19).

### MRI acquisition

The participants were scanned using the same 1.5 T scanner (Avanto, Siemens Medical Solutions; Erlangen, Germany) equipped with a 12-channel head coil. For rs-fMRI we used a T2* weighted echo-planar imaging (EPI) sequence (repetition time (TR) = 3000 milliseconds (ms), echo time (TE) = 70 ms, flip angle (FA) = 90°, voxel size = 3.44 × 3.44 × 4 millimetre (mm), field-of-view (FOV) = 220, descending acquisition, GeneRalized Autocalibrating Partial Acquisition (GRAPPA) acceleration factor = 2), 28 transversally oriented slices, no gap, with a scan time of 7 minutes and 30 seconds, yielding 150 volumes. Three dummy volumes were collected to avoid T1 saturation effects. Structural MRI data were collected using a three dimensional T1-weighted Magnetization Prepared Rapid Gradient Echo (MP-RAGE) sequence with the following parameters: TR / TE / time to inversion / FA = 2400 ms / 3.61 ms / 1000 ms / 8°, matrix 192 × 192, field of view = 240. Each scan lasted 7 minutes and 42 seconds and consisted of 160 sagittal slices with a voxel size of 1.20 × 1.25 × 1.25 mm.

### fMRI pre-processing and analysis

fMRI analysis was performed using FMRI Expert Analysis Tool (FEAT) Version 6.00, from FMRIB’s Software Library (20, 21). Head motion was corrected using MCFLIRT (22) before linear trends and low-frequency drifts were removed (high-pass filter of 0.01 Hertz). Image sequences were examined for excessive head motion causing image artefacts. FSL Brain extraction tool (23) was used to remove non-brain tissue. Spatial smoothing was performed using a Gaussian kernel filter with a full width at half maximum (FWHM) of 6 mm (24). FMRIB’s Nonlinear Image Registration tool (FNIRT) was used to register the participants fMRI volumes to Montreal Neurological Institute (MNI) 152 standard template using the T1-weighted scan as an intermediate, which had the non-brain tissue removed using procedures for automated volumetric segmentation in Freesurfer 5.3 (http://surfer.nmr.mgh.harvard.edu/) (25).

Single-session independent component analysis (ICA) was performed for all runs using Multivariate Exploratory Linear Optimized Decomposition into Independent Components (MELODIC) (26). The single-session ICA were submitted to FIX (27) for automatic classification into signal and noise components, in order to remove noise components from fMRI data. Data cleaning also included correction based on the estimated motion parameters for each run, using linear regression. FIX has been shown to effectively reduce motion induced variability, outperforming methods based on regression of motion parameters or spikes in the dataset (28).

The cleaned and MNI-conformed rs-fMRI datasets were submitted to temporal concatenation group independent component analysis (gICA) using MELODIC (26) with a model order of 30. These group level spatial components were then used as spatial repressors against the original rs-fMRI datasets to estimate subject-specific components and associated time series (dual regression (29)). The second group ICA component, encompassing the regions of the canonical DMN including the PCC, angular gyrus and mPFC, was thresholded at z>4 and used as a mask for extracting the mean DMN connectivity value from the subject specific dual-regression maps (Fig 1). The threshold z>4 (p=0.00006) was pragmatically chosen based on previous experience.

**Fig 1.**
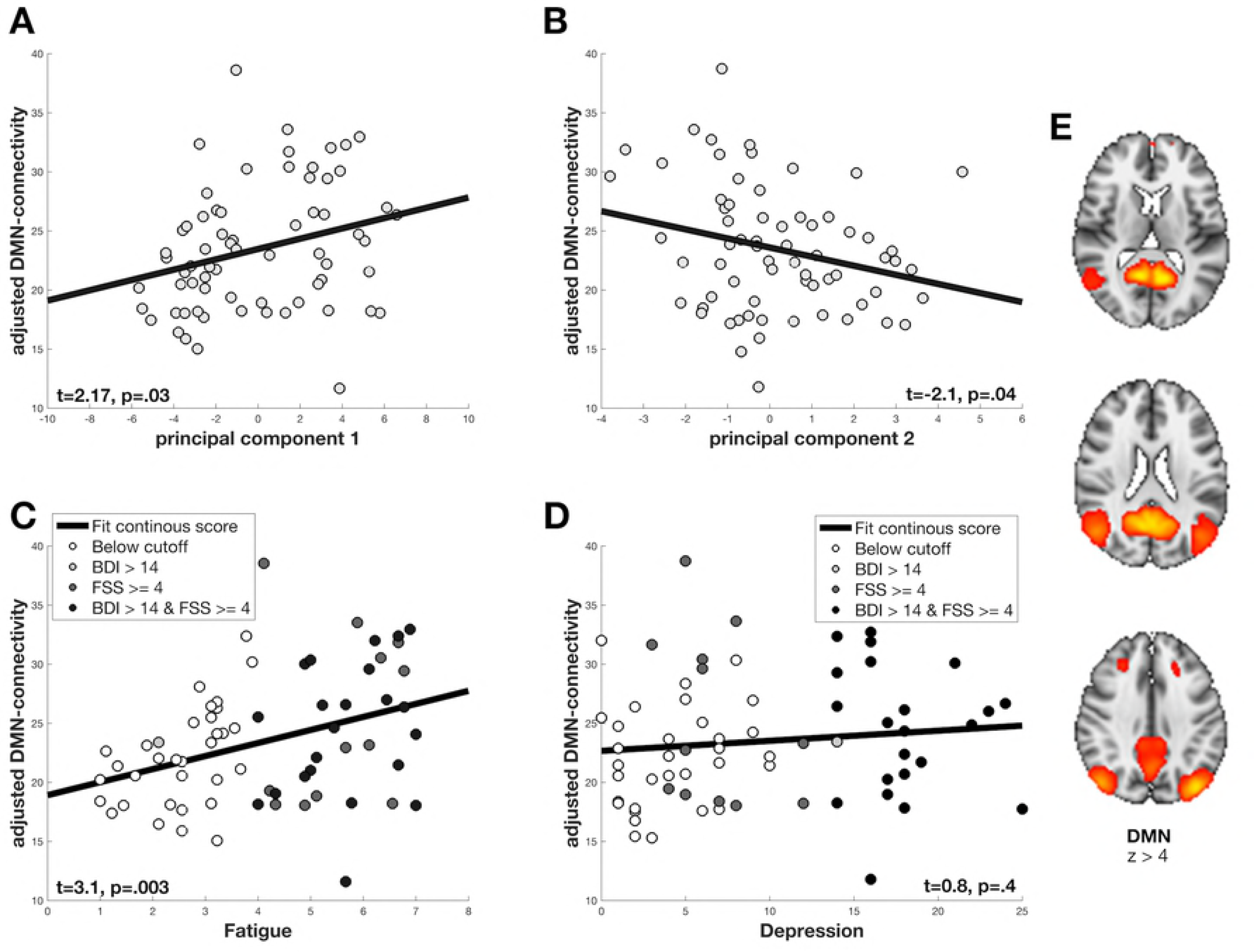
Associations between clinical symptoms and DMN connectivity. The correlation between adjusted DMN connectivity with the PCA components in A and B, and between adjusted DMN connectivity with FSS and BDI continuous scores in C and D. The grey tones for each subject represent clinical categories in C and D as described and shown in Table 1, and individual subject scores in A and B. (A) Increased PCA1 (high burden of both fatigue and depression) is positively correlated with DMN connectivity. (B) Decreased PCA2 (low burden of fatigue and high burden of depression) is negatively correlated with DMN connectivity. (C) Mean FSS correlated with DMN connectivity. (D) BDI sum scores correlated with DMN connectivity. Shown in E is the DMN component from the group independent component analysis (gICA). The component z-statistic map was thresholded at z>4. Depicted in three axial slices the posterior cingulate cortex (PCC) and the medial prefrontal cortex (mPFC) are masked out in red and yellow colours bilaterally.

### Statistical analyses

We used MATLAB version 9.2 (The MathWorks Inc., Natick, MA, 2017) and R (30) (R Core Team, Vienna, 2018) for statistical analyses. BDI and FSS subscores for all participants were submitted to PCA, decomposing the data into orthogonal components. To increase the statistical power of the PCA, we kept the seven MS patients missing fMRI data. The PCA yielded component loading coefficients for each questionnaire as well as component subject scores, resulting in a ranked list of PCA components with their associations to each BDI and FSS subscores (Fig 2). The subject scores for the two highest ranked PCA components were extracted for further analysis to test for associations with DMN connectivity.

**Fig 2.**
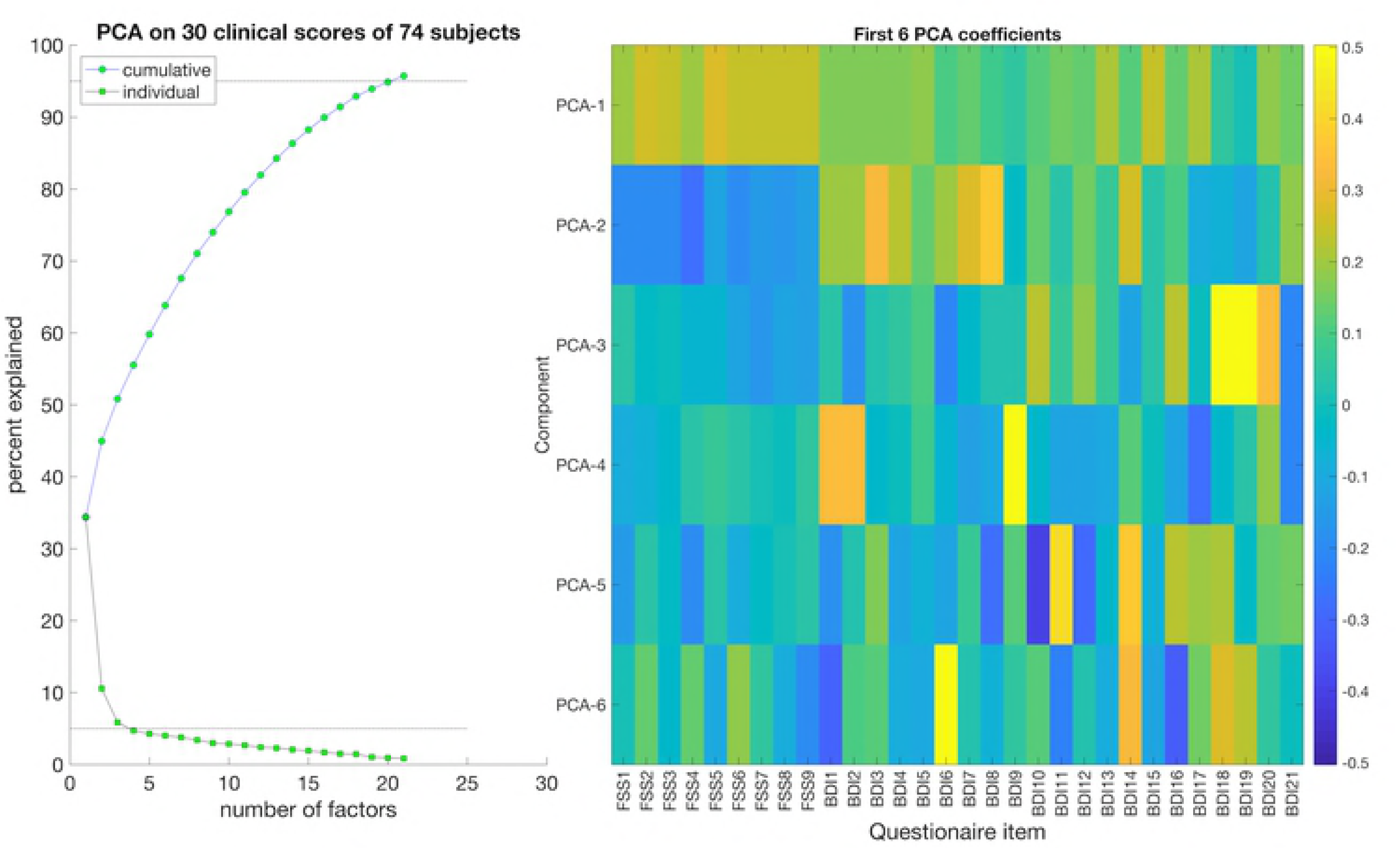
PCA from FSS and BDI subscores. PCA based on 30 clinical subscores (nine FSS and 21 BDI) for all participants. Left: The cumulative and individual explained variance of each PCA of the total variation in the clinical subscores. Right: A heatmap showing the first six PCA factors and their item loading on each component. Yellow and green boxes indicate association with high scores, while the blue boxes indicate association with low scores. The first PCA component (PCA1) captures common variance across BDI and FSS, while the second PCA component (PCA2) captures a pattern of covarying low FSS with high BDI scores.

Associations between DMN connectivity and clinical PCA scores were investigated using linear models, adjusting for age and sex. To evaluate effect sizes, we calculated Cohen´s f^2^, also taking into account age and sex. For Cohen´s f^2^ test, effect sizes are considered small (> 0.02), medium (> 0.15) and large (> 0.35). For clinical validation and comparison, we also estimated associations between DMN connectivity and the BDI and FSS continuous sum scores using multiple regression, adjusting for age and sex, and compared extreme groups based on conventional clinical thresholds (see above). To account for disability and cognitive impairment we also investigated the associations from the previously mentioned linear models with SDMT and EDSS scores.

## Data availability

Data cannot be shared publicly because of local restrictions for sensitive data.

## Results

### Participant demographics and characteristics

Table 1 summarizes demographic and clinical characteristics of the 74 included MS patients. The majority of the participants were women (70 %), mean age was 35.0 years (range 21-49 years). The majority of the participants received DMT, whereas 20 % of the participants were never treated. The participants were included on average 14.1 months after the date of diagnosis and disease duration was on average 73.0 months (range 5-272 months).

**Table 1.** **Demographic and clinical characteristics of the participants.**

| ***(a) Demographic characteristics*** | ***Patients (n = 74)*** |
| --- | --- |
| Female, n (%) | 52 (70) |
| Age, mean years (range) | 35.0 (21-49) |
| **Education** |  |
| Years, mean (range) | 14.9 (9-21) |
| ≥ 15 years education n (%) | 51 (69) |
| **Working status** |  |
| Unemployed or 100 % sick leave, n (%) | 7 (9) |
| Working (part- og full-time), student or maternity leave, n (%) | 67 (91) |
| ***(b) Clinical characteristics*** | ***Patients (n = 74)*** |
| **Neurological disability** |  |
| EDSS, mean (range) | 2.0 (0-6.0) |
| Number of total attacks, mean (range) | 1.8 (0-5) |
| **DMT** |  |
| No DMT, n (%) | 15 (20) |
| Active DMTs, n (%) | 48 (65) |
| Highly active DMTs, n (%) | 11 (15) |
| Months on treatment before study, mean (range) | 9.4 (0-34) |
| Months since diagnosis, mean (range) | 14.1 (1-34) |
| Disease duration, mean months (range) | 73.0 (5-272) |
| **Cognitive disability** |  |
| SDMT, mean (range) | 52.4 (30-80) |
| ***(c) Self-reported questionnaires*** | ***Patients (n = 74)*** |
| **FSS** |  |
| FSS, mean (standard deviation (SD)) | 4.2 (1.7) |
| Clinically significant fatigue (FSS mean ≥ 4), n (%) | 41 (55) |
| **BDI** |  |
| BDI sum, mean (SD) | 9.1 (6.7) |
| Clinically significant depressive symptoms (BDI sum $\geq 14$), n (%) | 23 (31) |
| **FSS and BDI status** |  |
| No fatigue (FSS mean $< 4$) and no depression (BDI sum $< 14$), n (%) | 32 (43) |
| Fatigue (FSS mean $\geq 4$) and no depression (BDI sum $< 14$), n (%) | 19 (26) |
| No fatigue (FSS mean $< 4$) and depression (BDI sum $\geq 14$), n (%) | 1 (1) |
| Fatigue (FSS mean $\geq 4$) and depression (BDI sum $\geq 14$), n (%) | 22 (30) |

EDSS, Expanded Disability Status Scale; DMT, disease modifying treatment; SDMT, symbol digits modalities test; FSS, Fatigue Severity Scale; BDI, Beck Depression Inventory

Fifty-five percent of all participants had clinically significant fatigue based on the FSS mean scores (FSS ≥ 4), and 31 % of all participants had clinically significant depressive symptoms based on BDI sum scores (BDI > 14). There were no significant differences in FSS and BDI scores between patients with and without rs-fMRI. The first PCA component (PCA1), which reflected common variance across depression and fatigue (high FSS and BDI scores), explained 34 % of the total variance in all FSS and BDI items (Fig 2). The second PCA component (PCA2), which reflected a characteristic pattern of low FSS with high BDI scores, explained 10 % of the total variance in all FSS and BDI subscores (Fig 2).

### Associations between clinical scores and DMN connectivity

Linear models revealed a significant positive correlation between PCA1 and DMN connectivity with small effect size (Cohen’s f^2^=0.075, t=2.17, *p*=0.03), indicating higher DMN connectivity with higher symptom burden. PCA2, which reflected a characteristic pattern of low FSS scores with high BDI scores, showed a significant negative correlation with DMN connectivity with small effect size (Cohen’s f^2^=0.067, t=-2.1, *p*=0.04) (Fig 1). Linear models revealed a significant positive correlation between FSS continuous mean scores correlated with DMN connectivity (t=3.1, *p*=0.003), and a non-significant positive association for BDI continuous sum scores correlated with DMN connectivity (t=0.8, *p*=0.39).

### Confounding effects of SDMT and EDSS

SDMT showed no significant association with DMN connectivity (t=1.7, p=0.09). The positive association between PCA1 and DMN connectivity remained significant (t=3.0, p=0.0045) when including SDMT in the model. The same model revealed a positive association between DMN connectivity and SDMT (t=2.6, p=0.011). The association between PCA2 and DMN became non-significant (t=-1.9, p=0.061) when including SDMT in the model. The same model revealed a non-significant positive association between DMN connectivity and SDMT (t=1.6, p=0.12).

EDSS showed no significant association with DMN connectivity (t=0.3, p=0.77). The positive association between PCA1 and DMN connectivity remained significant (t=2.2, p=0.031) when including EDSS in the model. The same model showed a non-significant association between DMN connectivity and EDSS (t=-0.51, p=0.61). The negative association between PCA2 and DMN connectivity remained significant (t=-2.0, p=0.049) when including EDSS in the model. The same model revealed a non-significant positive association between DMN connectivity and EDSS (t=0.25, p=0.81).

A linear model with FSS revealed a significant positive association with EDSS (t=2.5, p=0.014). The same model showed a non-significant association with SDMT (t=-1.0, p=0.32). A linear model with BDI revealed non-significant associations with EDSS (t=1.6, p=0.12) and SDMT (t=-1.4, p=0.17).

## Discussion

This study is to our knowledge among the first to study the complex interaction of fatigue and depression in patients with MS by multivariate decomposition analyses of these symptoms in relation to DMN connectivity measured by rs-fMRI. To understand the variability and mechanisms of both fatigue and depression is a key clinical question in MS.

Fatigue and depression represent common and strong predictors for quality of life in patients with MS, yet the pathophysiological mechanisms of fatigue and depression in MS patients are poorly understood. Converging lines of evidence have suggested associations between different symptoms (such as fatigue, cognitive impairment, depression) and the organization and synchronization of large-scale brain networks as measured by fMRI (3). Here, using multivariate decomposition of symptoms scores and rs-fMRI data we report significant association between DMN connectivity and both common and unique symptoms of depression and fatigue in patients with MS. The symptoms presenting in MS patients vary between individuals and is assumed to result primarily from demyelination and microscopic CNS tissue damage (3). Structural MRI studies have found diverse patterns of cortical thickness to be associated with different MS symptoms (8). Our results show correlation between DMN functional connectivity and FSS and BDI scores in MS, which support and further adds to previous knowledge.

One third of the participants in our study had both fatigue and depression, in line with other studies of MS patients (8). It is important to underline, that in this study, as in most MS papers, depressive symptoms are evaluated by self-reported psychometric scales, and no formal diagnosis of depressive mood disorder has been made (5). We found a significant positive correlation between DMN connectivity and the burden of fatigue and depression (PCA1 in Fig 1). DMN hyperconnectivity has been demonstrated in depression (31). A recent study investigated functional connectivity changes in MS patients with depression and suggested a functional link between depression and cognitive impairment (13). A functional link between depression and Alzheimer’s disease has also been reported (32). The same study proposed that depression in MS patients is a result of the demyelination and microscopic CNS tissue damage itself, and not a secondary symptom (13). A study on primary and secondary progressive MS patients found associations between cognitive impairment and reduction in resting state connectivity (33). Our findings support the hypothesis that symptoms of depression and fatigue are associated with altered DMN connectivity in MS, possibly influencing the normal function of the DMN as a critical hub of integration and flow of information.

We identified a second PCA component (PCA2) to be associated with a low burden of fatigue and a high burden of depressive symptoms. The second PCA component was negatively correlated with DMN connectivity, indicating that the clinical presentation of fatigue with no depression was associated with DMN hyperconnectivity. DMN hyperconnectivity in fatigue has been demonstrated in a group of breast cancer survivors, where enhanced intrinsic DMN connectivity with the frontal gyrus was associated with persistent fatigue after completed treatment (34). Our results indicate that there is hyperconnectivity in fatigued MS patients unrelated to depression, possibly caused by the inflammation or structural damage in the brain. Our findings of different DMN patterns depending on the symptom burden of fatigue and depression, may reflect the heterogeneity of symptoms in MS patients, reported in a recent review (4).

When adjusting our findings for cognitive impairment, the positive correlation of the first PCA component with DMN connectivity increased while the negative correlation with the second PCA component were slightly decreased. Disability did not have a confounding effect on the correlation between the PCA components and DMN connectivity. Yet we found a significant positive correlation between FSS and EDSS, in the way that higher disability was associated with higher symptoms of fatigue. Adjusting for cognitive impairment therefore seemed to only strengthen our results, while when adjusting for disability our results remained the same.

Our sample size is modest, but the participants were very thoroughly characterized and comprise a relatively homogenous group in terms of age, cognitive and physical disability, disease duration, education and clinical course. Concerning fatigue, the participants in our study scored a mean of 4.2 for FSS, which is lower than reported in some larger studies (35). However, the FSS scores for the participants included in this study were in line with a recent Norwegian MS study (6). Fatigue may impair the quality of life and contribute to the establishment and maintenance of depressive symptoms (4). The mean BDI sum score in our dataset was 9.1, which is lower than reported in some studies (5), but comparable with a Swedish study (36). Possible reasons for the relatively low BDI sum score in our sample include the low age, newly diagnosed RRMS, short disease duration and few brain lesions in our MS patients (15).

The structural underpinnings in the brain of the observed associations are not known, and future studies combining structural MRI and fMRI data could give further insights into the pathophysiology of depression and fatigue in MS. However, the associations between symptoms of fatigue and depression with DMN connectivity identified by rs-fMRI in our study suggest different pathophysiology for the two most prominent components revealed by the multivariate decomposition analysis of symptoms of fatigue and depression. Previous studies assessing cortical morphometry in an overlapping patient sample reported regional associations between cortical surface areas and several clinical manifestations, where the most prominent structural association were smaller cortical surface area and volume significantly associated with depressive symptoms (15).

In addition to our modest sample size, other limitations should be considered when interpreting our results. We did not include lesion filling as part of the fMRI analysis pipeline, as this was not implemented. In MS patients, permanent damage affects the white matter of the CNS and can cause disconnection syndromes (3). The functional connectivity and large-scale networks depend on structural connections, and inter-individual variability in DMN connectivity, and its association with clinical traits, might be mediated by degree of demyelination, atrophy of both the grey and white matter and microscopic CNS damage (12). We did not have access to a healthy control sample. Yet, our results only focus on the DMN connectivity changes in relation to neuropsychiatric symptoms within the MS group. Future studies are needed to test if our results can be generalized to other populations.

## Conclusion

In conclusion, multivariate decomposition of FSS and BDI symptom data supported that the clinical manifestations of fatigue and depression in patients with MS reflect both overlapping and unique variability in the FSS and BDI subscores. The observed differential correlations between symptoms of fatigue and depression and DMN connectivity underline the heterogeneity and complexity of fatigue and depression in MS. Our analyses revealed that high burden of both fatigue and depression was associated with DMN hyperconnectivity, while we also found hyperconnectivity in DMN to be associated with high burden of fatigue in absence of depression. Effect sizes were in general relatively small, and further investigations into the mechanisms of fatigue and depression in MS are warranted. Multivariate decomposition analyses of MS symptoms in relation to default mode network (DMN) connectivity measured by resting-state-fMRI (rs-fMRI) is a promising method to pursue these questions.

## Acknowledgment and funding

We thank all the patients who participated in our study. The project was supported by grants from The Research Council of Norway (240102,) and the South-Eastern Norway Regional Health Authority (2011059/ ES563338/Biotek 2021, 2014097) (Principal Investigator for both grants: HFH). The funders had no role in study design, data collection and analysis, decision to publish or preparation of the manuscript.

